# A Visualization Tool to Evaluate Pairwise Protein Structure Alignment Algorithms

**DOI:** 10.1101/342899

**Authors:** Shalini Bhattacharjee, Asish Mukhopadhyay

**Affiliations:** School of Computer Science, University of Windsor, Ontario, Canada; Toronto Star, Toronto, Canada

**Keywords:** Protein Structure Alignment, Principal Component Analysis, Clustering, Algorithms

## Abstract

The alignment of two protein structures is a fundamental problem in structural bioinformatics. In this paper, we propose a novel approach to measure the effectiveness of a sample of three such algorithms, *DALI, TM-align* and *EDAlign_sse_*. The underlying premise of our approach is that structural proximity should translate into spatial proximity.

## 1 Introduction

A protein molecule is a linear polypeptide chain, with adjacent pairs of amino acids joined together by peptide bonds, giving rise to the nomenclature “polypeptide”. In order to perform its particular biological function, the linear polypeptide chain folds into a stable, low-energy 3-dimensional tertiary structure. The latter structure is formed by loops joining together two types of secondary structures, known as *α*-helices and *β*-sheets.

Since the functionalities of a protein are determined by their 3d-structures, it became an important task to determine the functionalities of a newly sequenced protein by comparing its structure with the structures of proteins of known functionalities. Thus algorithms for determining the structural similarities of proteins became a very active area of research, giving birth to many popular algorithm like DALI, TM-align, CE, Structal, to name a few. As Godzik observed [4], determining the structural similarity of two proteins is not an easy problem. Among the major hurdles are the problems of choosing a suitable structural abstraction of a protein molecule, crafting an appropriate measure of similarity, and finding a set of equivalent (or corresponding) pairs that helps in finding a rigid transformation for the actual alignment by minimizing an error function.

The alignment of two protein structures is the 3-dimensional analogue of linear sequence alignment of peptide or nucleotide sequences. An initial equivalence set can be obtained by various methods such as comparison of distance matrices [8], maximal common subgraph detection, geometric hashing, local geometry matching, spectral matching, contact map overlap and dynamic programming. This equivalence set is optimized by different methods such as a Monte Carlo algorithm or simulated annealing, dynamic programming, incremental combinatorial extension of the optimal path [17] and genetic algorithm. Indeed the goal is to determine an alignment of protein residues to measure the extent of structural similarity. To quantify this similarity, various measures have been defined and can be broadly classified into four categories: (1) distance map similarity (2) root mean square deviation (*RMSD*) (3) contact map overlap [5] (4) universal similarity matrix. A comprehensive list of different similarity measures are discussed by Hasegawa and Holm [6]. Surprisingly, even after so many years of research there is no universally acknowledged definition of similarity score to measure the extent of structural similarity.

In a seminal work Godzik [4] compared several well-known structural alignments algorithms with different similarity measures, each focusing on a different aspect of structural similarity, with respect to three different pairs of proteins and showed that this problem has no unique answer. In this paper, we propose a novel approach to measure the effectiveness of four structural alignment algorithms, *DALI, TM-align, EDAlign_sse_* and *CE.* The underlying premise of our approach is that structural proximity should translate into spatial proximity. To verify this, we carried out extensive experiments with five different datasets, each consisting of proteins from two to six different families. For each dataset, we computed a distance matrix, where each distance is the *cRMSD* distance of a pair of proteins. For each distance matrix, we have used Principal Component Analysis to obtain an embedding of a set of points (each representing a protein) that realize these distances in a two-dimensional space. To compare the clustering of the families, we have used the *k*-means clustering algorithm to cluster the points, sans family labels. Our conclusion is that of the three algorithms, *TM*-align is most successful in translating structural proximity to spatial proximity.

## 2 Notations and Definitions

A protein is modeled as a sequence of points, *P* = {*p_i_*|*p_i_* ∊ *R*^3^, *i* = 1, 2, 3, …, *m*}, in a 3-dimensional Euclidean space, where *m*(= |*P*|) is the number residues and *p_i_* represents the coordinates of the central *α*-carbon atom of the *i*-th residue. In what follows, we will use this sequence *P* to refer to the protein it represents.

Given two proteins *P* and *Q* of length *m* and *n* respectively, an alignment of *P* and *Q* is a sequence of corresponding pairs of points of *P* and *Q,*

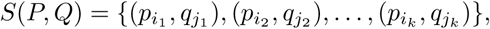

where 1 ≤ *i*_1_ < *i*_2_ < … < *i_k_* ≤ *m* and 1 ≤ *j*_1_ ≠ *j*_2_ ≠ … ≠ *j_k_* ≤ *n,* together with a rigid transformation 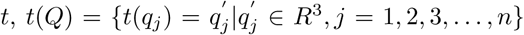, that optimizes some similarity measure for the above correspondence.

### 2.1 Similarity measures

To measure the extent of structural similarity of two proteins, the root mean square deviation (*RMSD*) is widely used [10, 11]. Two different *RMSD* measures have been proposed in the literature: (1) coordinate root mean square deviation (*cRMSD*) and (2) distance root mean square deviation (*dRMSD*). For two aligned substructures of proteins *P* and *Q* of length *k,* the *cRMSD* measure is defined below.
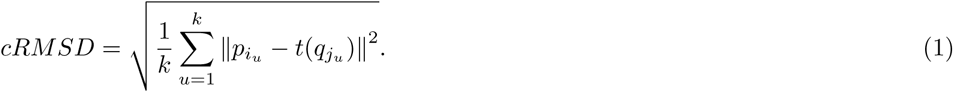

Since the similarity measures, *cRMSD* and *dRMSD,* are in terms of absolute distances, a small presence of outliers may result in a poor *RMSD* even if the two structures are globally similar. To circumvent this problem, Zhang and Skolnick [19] introduced a sequence independent structural alignment measure (TM-score) that is a variation of a measure originally defined by Levitt and Gerstein [12]. A critical assessment of this TM-score has been given by Xu and Zhang [18].

Given two proteins, a template protein *P* and a target protein *Q*, |*P*| ≥ |*Q*|, the structural similarity is obtained by a spatial superposition of *P* and *Q* that maximizes the following score
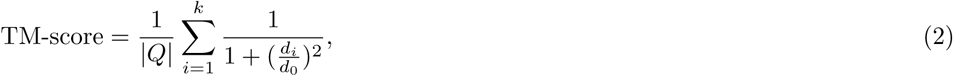

where *k* is the number of aligned residues of *P* and *Q*; *d_i_* is the distance between *i*-th pair of aligned residues and 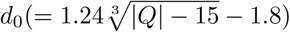 is a normalization factor.

When the value of *d*_0_ in equation (2) is set to 5*A^o^*, the resulting TM-score is known as a raw TM-score (rTM-score). Xu and Zhang [18] observed that two proteins are structurally similar and belong to the same fold when the TM-score > 0.5.

## 3 Materials and Methods

### 3.1 Creating Datasets

We have selected protein families which are completely different from one another structurally, implying divergence in functional behavior. The proteins that make up our datasets have been drawn from the following families.

`Myosins`: a class of motor proteins that are crucial to muscle contraction and other motility processes in eukaryotes.
`GTPases`: a large family of hydrolase enzymes that bind and hydrolase guanosine triphosphate (GTP). These help in regulating cell growth, cell differentiation and cell migration [1].
`Caspases`: a family of protease enzymes playing essential roles in programmed cell death and inflammation [2].
`EF Hand Proteins`: a large family of calcium-binding proteins, each with an EF-hand or alpha-loop-alpha motif [13].
`Calmodulin` is a calcium transducer. It is a calcium-binding protein that can bind to and regulate the functions of different protein targets, thereby affecting many different cellular functions [14].
`Phosphotransferase`: a class of enzymes that catalyze phosphorylation reactions. Phosphorylation is crucial for protein function as it activates or deactivates nearly half of the enzymes. It is also a frequently-occurring post-translation modification in eukaryotic cells (source: Wikipedia).
`Cyclophilins`: a family of proteins found in vertebrates and other organisms that bind to cyclosporin A, an immunosuppressant commonly used to suppress rejection after an internal organ transplant [3].

The five datasets below were created from the above protein classes.

1. Two families, each consisting of 10 proteins

2. Three families, each consisting of 10 proteins

| Family No. | Family Name | Proteins (PDBs) |
| --- | --- | --- |
| 1 | GTPases | 1YZQ, 1T91, 1YR9, 1XTS, 1MKY, 1PUJ, 2RCN, 2MSC, 2X2E, 1CEE |
| 2 | Myosins | 1W7J, 1W7I, 1B7T, 1OE9, 2AKA, 2MYS, 4P7H, 2EC6, 2OS8, 2OGT |

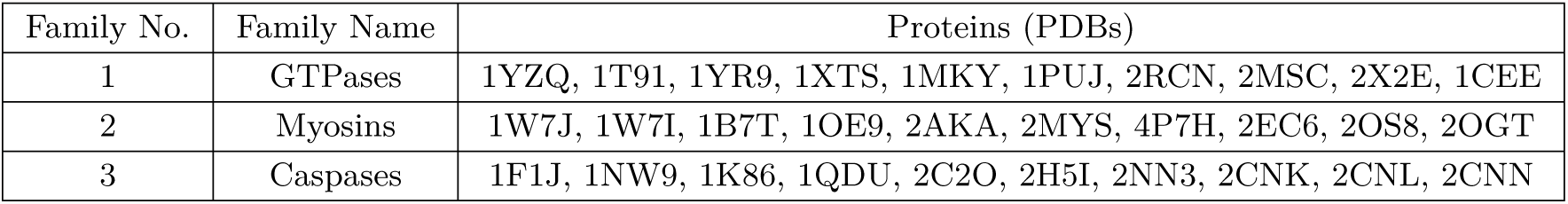
3. Four families, each consisting of 5 proteins

2. Three families, each consisting of 10 proteins
| Family No. | Family Name | Proteins (PDBs) |
| --- | --- | --- |
| 1 | GTPases | 1YZQ, 1T91, 1YR9, 1XTS, 1MKY, 1PUJ, 2RCN, 2MSC, 2X2E, 1CEE |
| 2 | Myosins | 1W7J, 1W7I, 1B7T, 1OE9, 2AKA, 2MYS, 4P7H, 2EC6, 2OS8, 2OGT |
| 3 | Caspases | 1F1J, 1NW9, 1K86, 1QDU, 2C2O, 2H5I, 2NN3, 2CNK, 2CNL, 2CNN |

4. Five families, each consisting of 6 proteins

3. Four families, each consisting of 5 proteins
| Family No. | Family Name | Proteins (PDBs) |
| --- | --- | --- |
| 1 | Caspases | 3DEI, 2C2O, 1K86, 2H5I, 1QDU |
| 2 | Myosins | 2AKA, 2EC6, 1OE9, 1B7T, 4P7H |
| 3 | ER Hand Proteins | 1XO5, 1IJ5, 2BE4, 2KQ6, 5BFX |
| 4 | Calmodulin | 1CLL, 1CFC, 3CLN, 1MXE, 1DMO |

5. Six families, each consisting of 7 proteins

4. Five families, each consisting of 6 proteins
| Family No. | Family Name | Proteins (PDBs) |
| --- | --- | --- |
| 1 | Caspases | 3DEI, 2C2O, 1K86, 2H5I, 1QDU, 1RHK |
| 2 | Myosins | 2AKA, 2EC6, 1OE9, 1B7T, 4P7H, 1W7I |
| 3 | ER Hand Proteins | 1XO5, 1IJ5, 2BE4, 2KQ6, 5BFX, 1JUO |
| 4 | Calmodulin | 1CLL, 1CFC, 3CLN, 1MXE, 1DMO, 2F2P |
| 5 | Phosphotransferase | 2B0G, 1FYN, 2HWG, 1J7U, 1ND4, 4ORK |

5. Six families, each consisting of 7 proteins
| Family No. | Family Name | Proteins (PDBs) |
| --- | --- | --- |
| 1 | Caspases | 3DEI, 2C2O, 1K86, 2H5I, 1QDU, 1RHK, 1V0D |
| 2 | Myosins | 2AKA, 2EC6, 1OE9, 1B7T, 4P7H, 1W7I, 1BR4 |
| 3 | ER Hand Proteins | 1XO5, 1IJ5, 2BE4, 2KQ6, 5BFX, 1JUO, 1JFK |
| 4 | Calmodulin | 1CLL, 1CFC, 3CLN, 1MXE, 1DMO, 2F2P, 2K0F |
| 5 | Phosphotransferase | 2B0G, 1FYN, 2HWG, 1J7U, 1ND4, 4ORK, 1PNJ |
| 6 | Cyclophilins | 1CWA, 1M9C, 1AK4, 2CPL, 2RMB, 2Z6W, 1NMK |

For each dataset, we ran the three pairwise alignment algorithms, *DALI* [9], *TM-align* [**?**] and *SSEAlign* [15], on each pair of proteins in the set to create a distance matrix. The distances are the *cRMSD* values of all the pairs. For example, from the first dataset, consisting of two families of ten proteins each, for a total of 20 proteins, we obtained a 20 × 20 symmetric distance matrix with 190 entries above the main diagonal.

### 3.2 Principal Component Analysis

Principal Component Analysis [7] is a dimensionality reduction technique that makes it possible to visualize data in low-dimensional spaces. In this method, a new set of variables are obtained from the existing set of variables by a linear combination of the existing variables. The new set of variables are called Principal Components. The components are obtained in such a manner that the first one accumulates the maximum variation of the existing data. The succeeding component have the next highest variation and so on. In whatever reduced dimension the existing data is embedded, this process preserves the highest possible variance of the original set of data [16].

### 3.3 *k*-means Clustering

The *k*-means clustering algorithm partitions a set of *n* data points in an *m*-dimensional Euclidean space into *k*-clusters. Each cluster consists of data points closest to the cluster center. The parameter *k* is part of the input to the clustering algorithm.

The data points are obtained by applying Principal Component Analysis to the *cRMSD* distance matrix. The returned set of points (representing proteins) lie in a low dimensional space (we have chosen 2 as the embedding dimension). The visualization of these points with their family labels show a natural clustering that demonstrates how well an alignment algorithm translates structural proximity to spatial proximity.

When the same set of points, sans family labels, are subjected to the *k*-means clustering algorithm which uses only the spatial proximities of the points, we expect the clustering to remain largely unchanged with respect to the natural clustering. We discuss how well the three alignment algorithms have fared with respect to our expectations in the section on experimental results.

### 3.4 Proposed Algorithm

We call our algorithm *EPAA,* short for Evaluating Pairwise Alignment Algorithms. Below, we provide a formal description of *EPAA* in pseudocode form.

#### Algorithm 1 EPAA

1: **for** Each alignment algorithm, 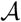 **do**

2: **for** Each dataset, 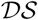 **do**

3: Run 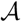 for every pair of proteins to build a *cRMSD* distance matrix

4: Input the distance matrix to the PCA algorithm to obtain a two dimensional embedding

5: Run the *k*-means clustering method on the embedded point set

6: Plot the points with original family labels

7: Plot the clusters obtained from the *k*-means algorithm, sans family labels

8: **end for**

9: **end for**

## 4 Experimental Results and Analysis

Dataset 1- Two families, each consisting of 10 proteins.

Referring to Figs.1-3, we find that with *TM-align,* the structural clustering (Fig. 2(a)) and the spatial clustering (Fig. 2(b)) are quite similar. The same cannot be said of *DALI* (Fig. 1) and *SSEAlign* (Fig. 3). In each of those cases, the proteins do not form two clearly distinguishable spatial clusters.

**Figure 1.**
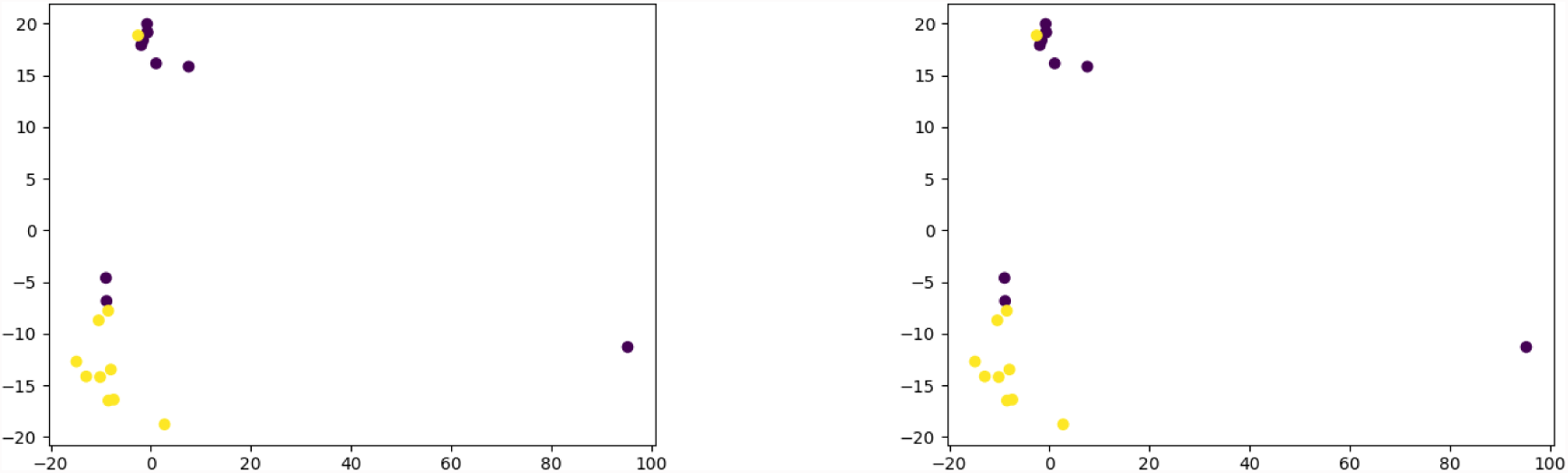
Clusters formed by *DALI*: (left) Known family labels (right) Unknown family labels

**Figure 2.**
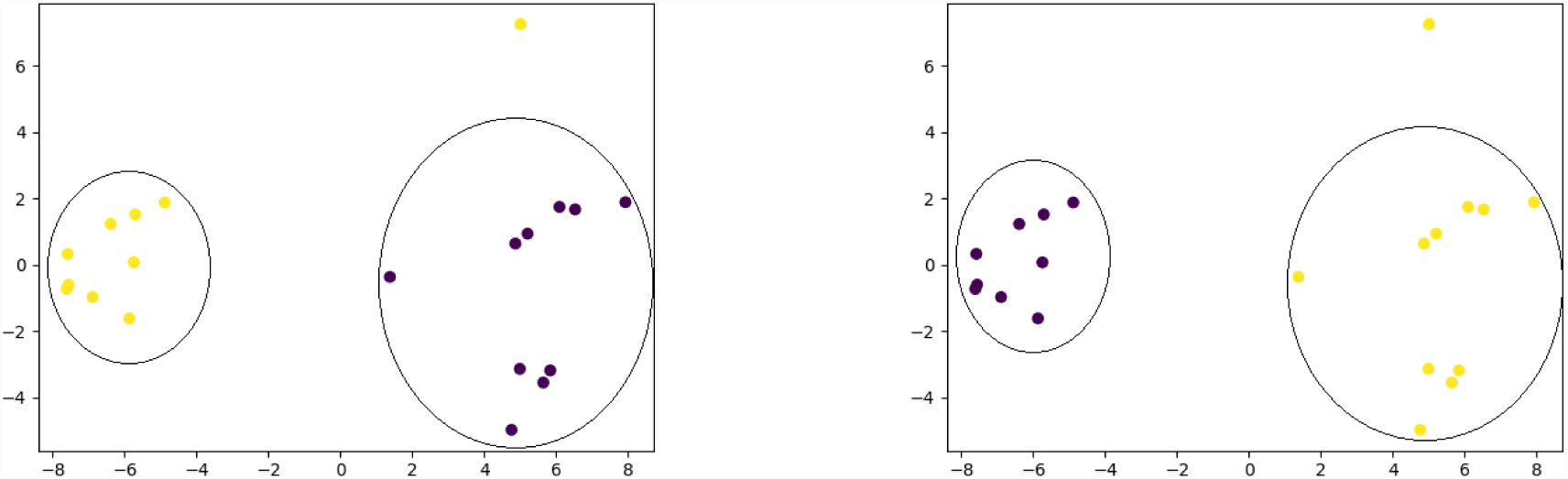
Clusters formed by *TM-align*: (left) Known family labels (right) Unknown family labels

**Figure 3.**
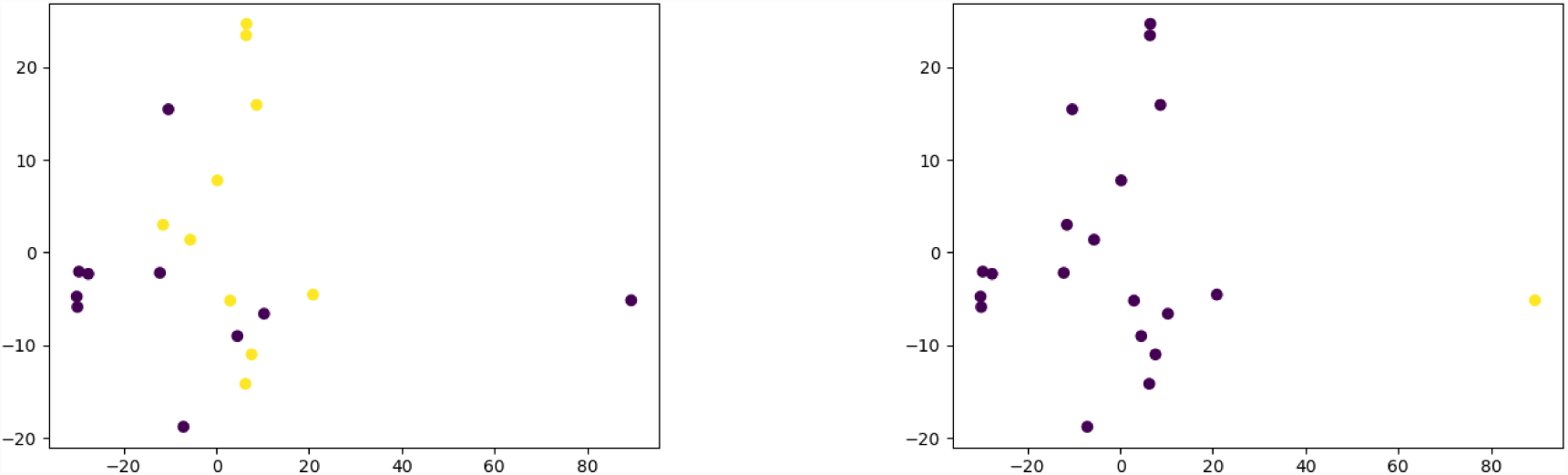
Clusters formed by EDAlign SSE: (left) Known family labels (right) Unknown family labels

Dataset 2- Three families, each consisting of 10 proteins

Referring to Figs. 4-6, we find that with *DALI* (Fig. 4) and *SSEAlign* (Fig. 6), the structural clustering (Fig. 4(a) and Fig. 6(a), respectively) and the spatial clustering (Fig. 4(b) and Fig. 6(b) respectively) are very different. For both of them, two families out of three got mixed up in the spatial clustering. In the case of *TM-align,* the structural clustering (Fig. 5(a)) and the spatial clustering (Fig. 5(b)) are almost the same. All the three families can be easily distinguished from the clusters formed.

**Figure 4.**
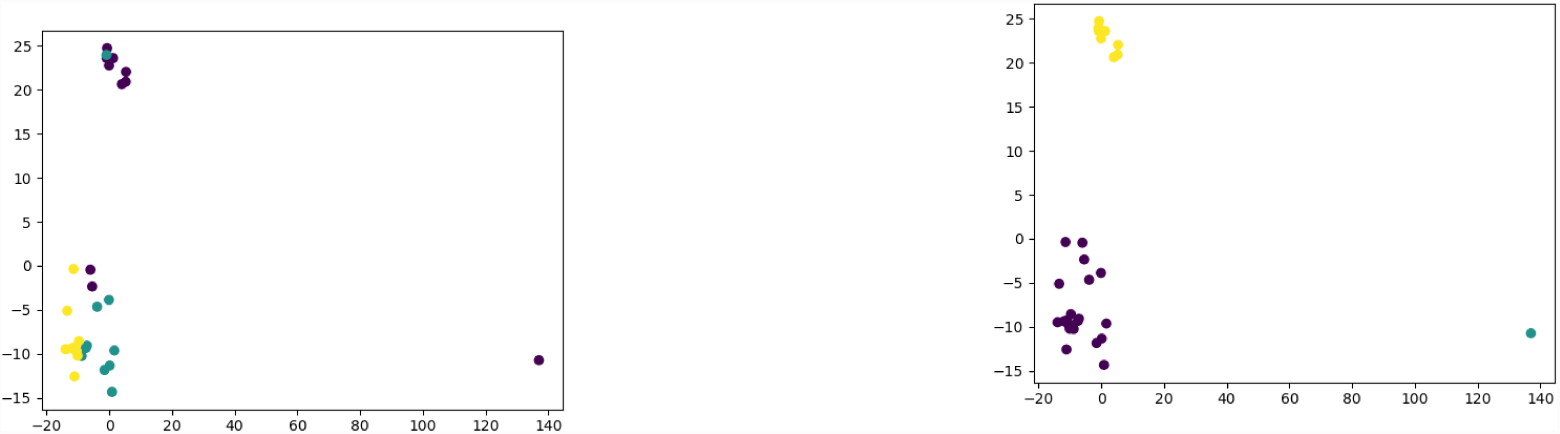
Clusters formed by *DALI*: (left) Known family labels (right) Unknown family labels

**Figure 5.**
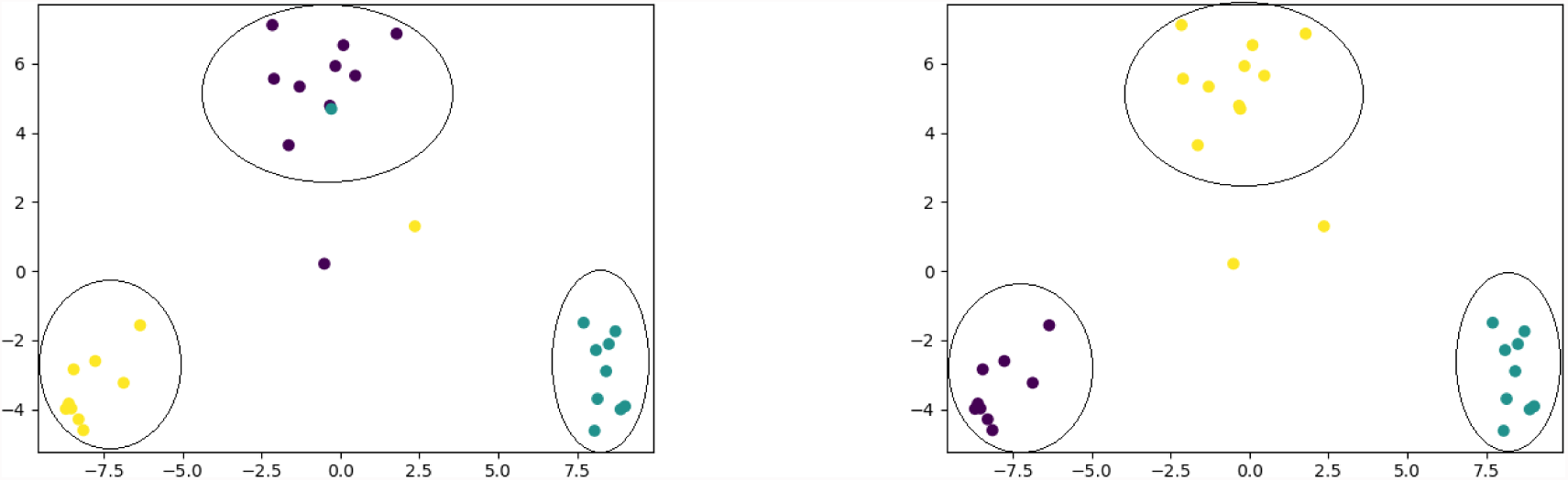
Clusters formed by *TM-align*: (left) Known family labels (right) Unknown family labels

**Figure 6.**
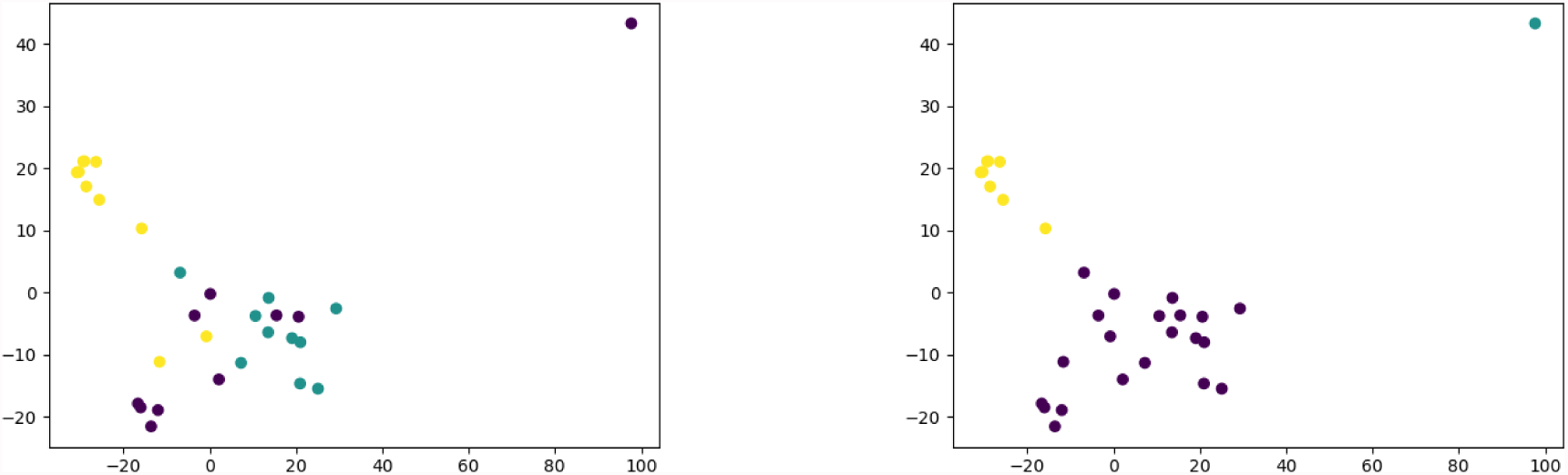
Clusters formed by EDAlign SSE: (left) Known family labels (right) Unknown family labels

Dataset 3- Four families, each consisting of 5 proteins

Referring to Figs. 7-9, we find that with *TM-align,* the structural clustering (Fig. 8(a)) and the spatial clustering (Fig. 8(b)) of the 3 families are nearly similar. But in case of *DALI* (Fig. 7), the spatial clustering of nearly all the families are dispersed. From Figs. 9(a) and 9(b), we can say that the clustering in *SSEAlign* is better than that of *DALI* as it has formed proper spatial clustering in two families out of four. However, it is not as good as *TM-align.*

**Figure 7.**
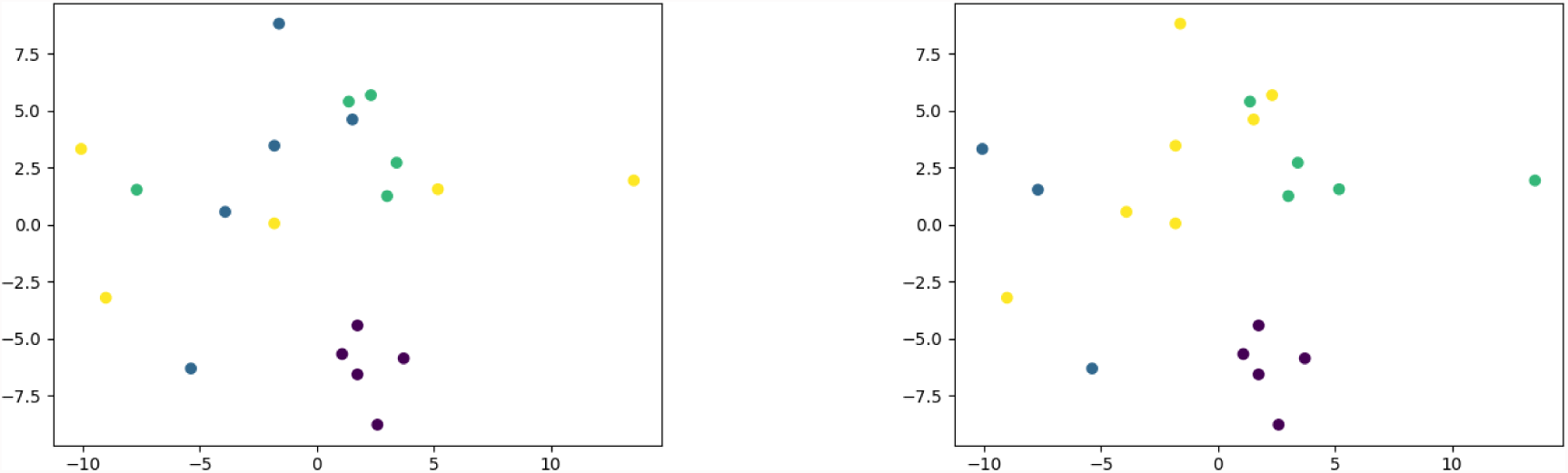
Clusters formed by *DALI*: (left) Known family labels (right) Unknown family labels

**Figure 8.**
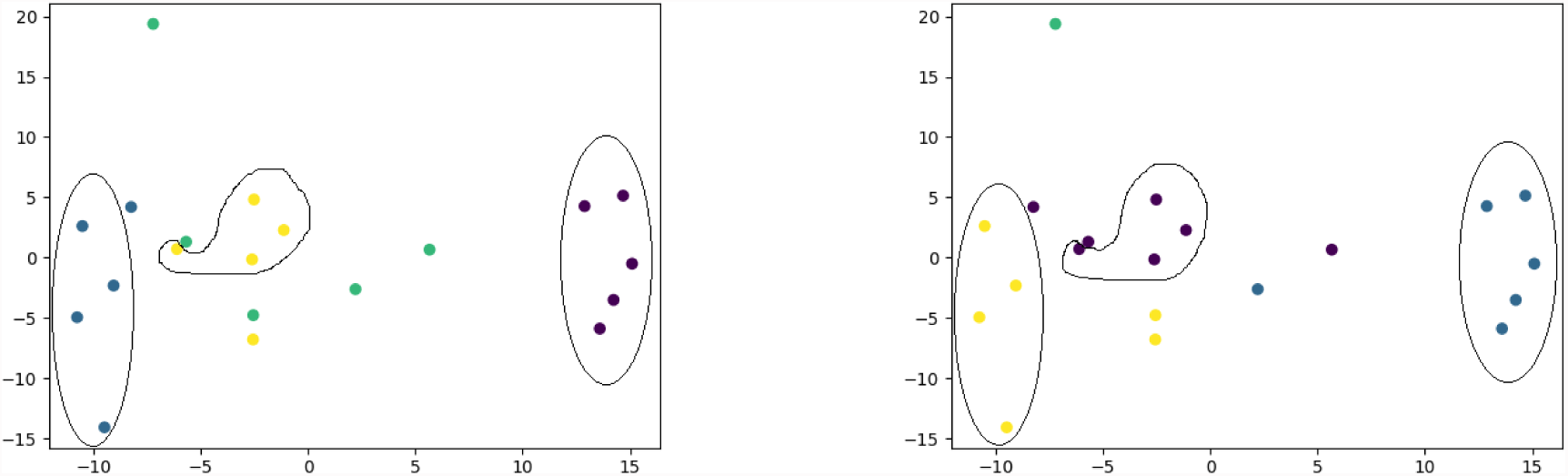
Clusters formed by *TM-align*: (left) Known family labels (right) Unknown family labels

**Figure 9.**
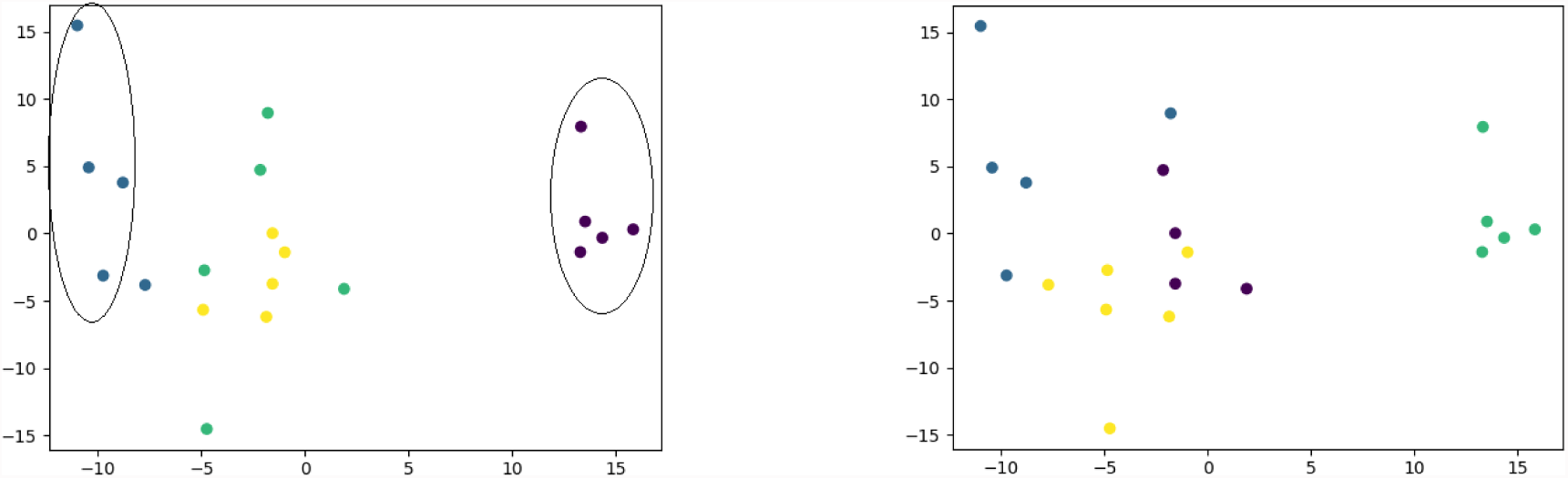
Clusters formed by EDAlign SSE: (left) Known family labels (right) Unknown family labels

Dataset 4- Five families, each consisting of 6 proteins

Referring to Figs. 10-12, we again find that for *TM-align* the structural clustering (Fig. 11(a)) and the spatial clustering (Fig. 11(b)) are similar for two to three families out of 5. Unlike *DALI* (Fig. 10) and *SSEAlign* (Fig. 12), *TM-align* has successfully formed a nice clustering even with the huge dataset of 30 proteins. We see that *SSEAlign* has got one family clustered properly, as compared to *DALI* which has failed for this large dataset.

**Figure 10.**
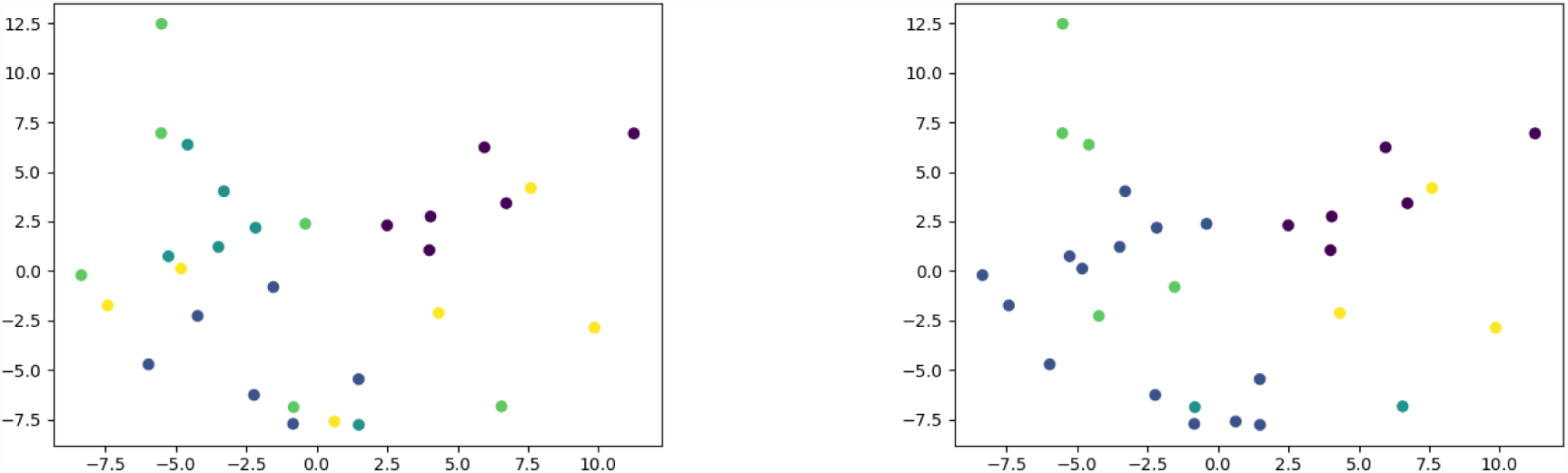
Clusters formed by *DALI*: (left) Known family labels (right) Unknown family labels

**Figure 11.**
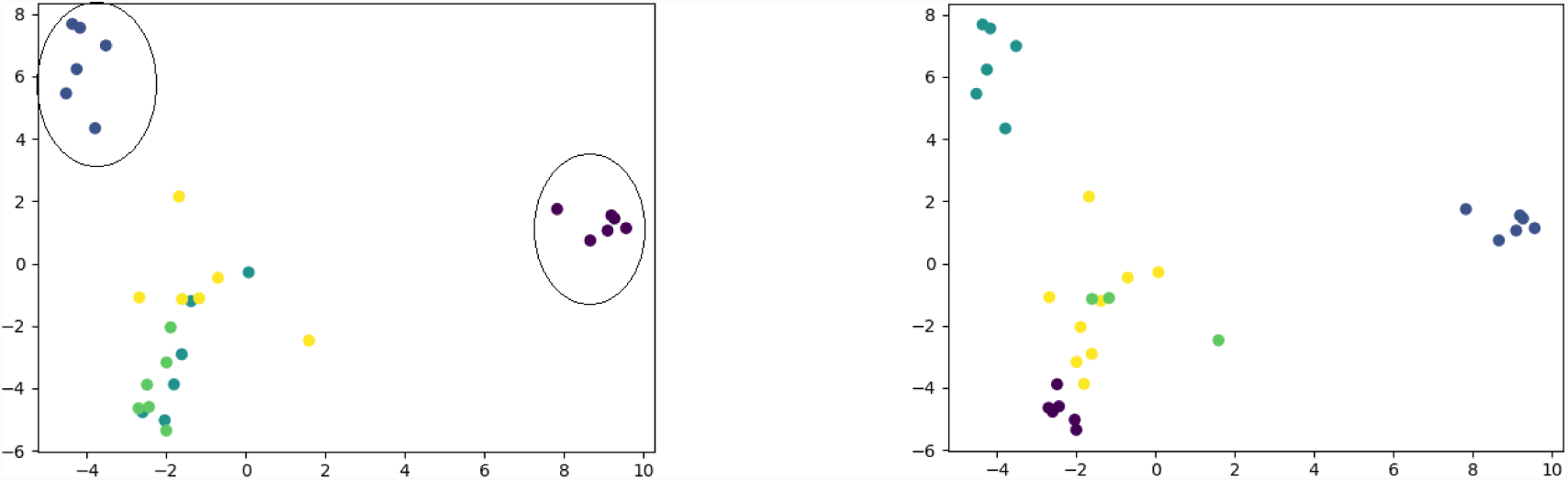
Clusters formed by *TM-align*: (left) Known family labels (right) Unknown family labels

**Figure 12.**
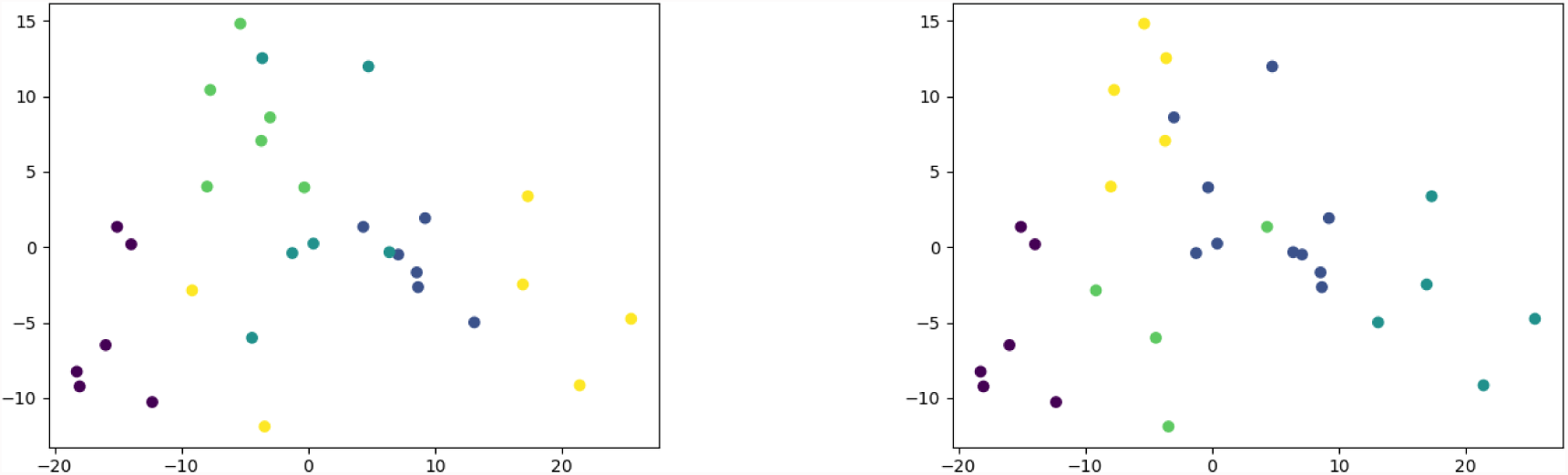
Clusters formed by EDAlign SSE: (left) Known family labels (right) Unknown family labels

Dataset 5- Six families, each consisting of 7 proteins

Referring to Figs. 13-15, we find that for this dataset *SSEAlign* has performed better than *TM-align.* The structural clustering (Fig. 15(a)) and the spatial clustering (Fig. 15(b)) of *SSEAlign* are similar for four families unlike *TM-align* which has similar clustering for 3 families. Most of the families are easily distinguishable. From Fig. 15(a) and Fig. 15(b) we find that the structural clustering and the spatial clustering for *DALI* again proves to be the worst with the proteins from different families all dispersed among the different clusters.

**Figure 13.**
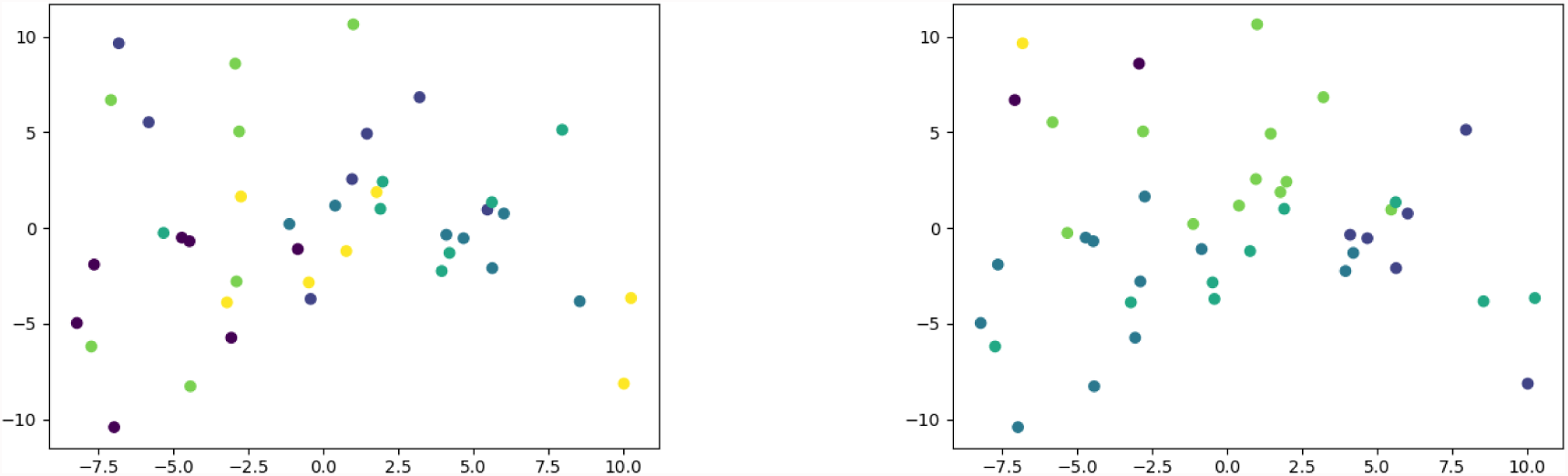
Clusters formed by *DALI*: (left) Known family labels (right) Unknown family labels

**Figure 14.**
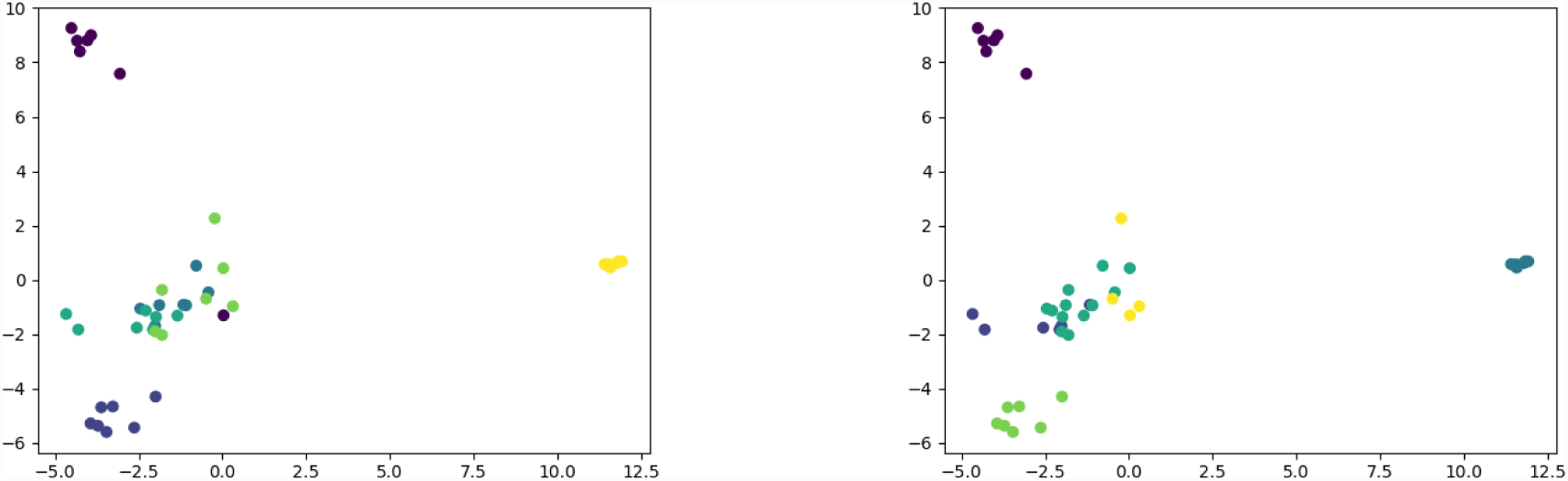
Clusters formed by *TM-align*: (left) Known family labels (right) Unknown family labels

**Figure 15.**
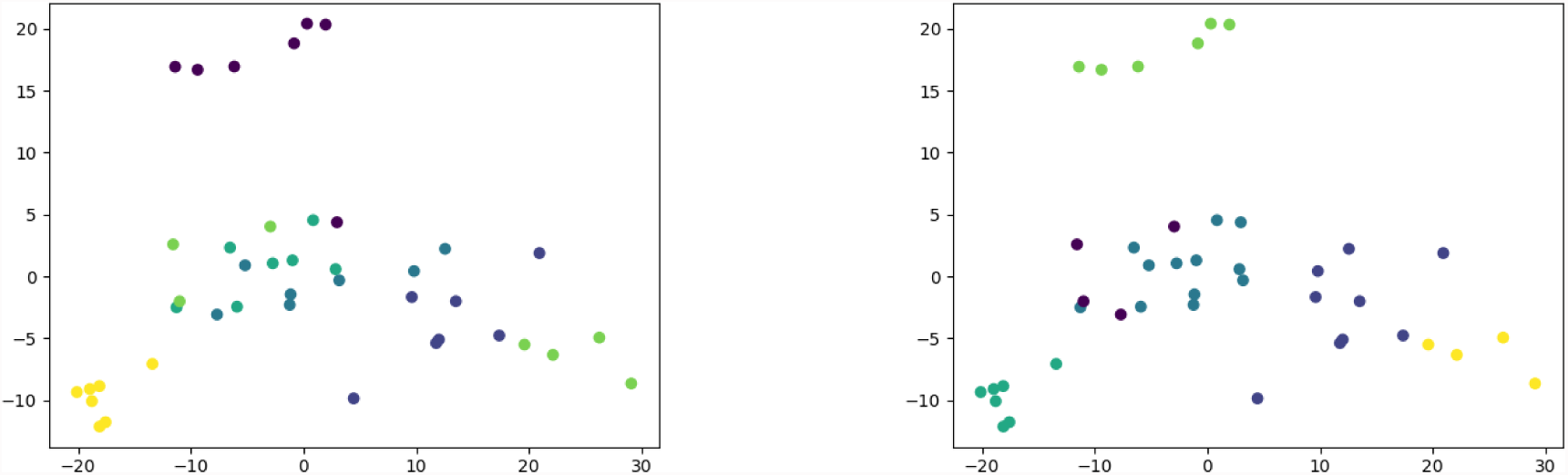
Clusters formed by EDAlign SSE: (left) Known family labels (right) Unknown family labels

## 5 Conclusions

The evidence from the experiments clearly suggest that of the three algorithms *TM-align* is the most successful one in correlating structural proximity to spatial proximity. In *TM-align,* the majority of the proteins were clustered according to their family, and there was minimal mix-up of the families as compared to the other two pairwise alignment algorithms. As for *EDAlignSSE,* it performed creditably as compared to *DALI*[9]; in fact, it managed to get some of the proteins clustered according to their families. This is particularly evident for dataset 3, in which case it formed 4 family clusters with similar proteins. In the final datasets, with 42 different proteins *DALI*[9] showed very poor clustering. The huge data set accentuated its weakness more emphatically. Therefore, based on our clustering technique to measure the effectiveness of an alignment algorithm, we conclude that *TM-align* is a better alignment algorithm than *EDAlignSSE* or *DALI* [9]. This work can be extended further to test and evaluate many other pairwise alignment algorithms proposed in the past. It can be used as a measure to find the quality of new pairwise alignment algorithms. This work can be also be used to infer evolutionary relationships from the clusters.

## References

1 Bruce Alberts, Dennis Bray, Karen Hopkin, Alexander Johnson, Julian Lewis, Martin Raff, Keith Roberts, and Peter Walter. Essential cell biology. Garland Science, 2013.

2 Howard Y Chang and Xiaolu Yang. Proteases for cell suicide: functions and regulation of caspases. Microbiology and molecular biology reviews, 64(4):821–846, 2000.

3 S Jane Flint, Vincent R Racaniello, Lynn W Enquist, Anna M Skalka, et al. Principles of virology, Volume 2: pathogenesis and control. Number Ed. 3. ASM press, 2009.

4 Adam Godzik. The structural alignment between two proteins: Is there a unique answer ? Protein Science, 5:1325–1338, 1996.

5 Adam Godzik, Andrzej Kolinski, and Jeffrey Skolnick. Topology fingerprint approach to the inverse protein folding problem. Journal of Molecular Biology, 227(1):227–238, 1992. URL:http://www.sciencedirect.com/science/article/pii/002228369290693E, doi: http://dx.doi.org/10.1016/0022-2836(92)90693-E.

6 Hitomi Hasegawa and Liisa Holm. Advances and pitfalls of protein structural alignment. Current Opinion in Structural Biology, 19(3):341–348, 2009. URL: http://www.sciencedirect.com/science/article/pii/S0959440X09000621, doi:http://dx.doi.org/10.1016/j.sbi.2009.04.003.

7 Steven M Holland. Principal components analysis (pca). Department of Geology, University of Georgia, Athens, GA, pages 30602–2501, 2008.

8 Liisa Holm and Chris Sander. Protein Structure Comparison by Alignment of Distance Matrices. Journal of Molecular Biology, 233(1):123–138, September 1993. URL: http://dx.doi.org/10.1006/jmbi.1993.1489, doi:10.1006/jmbi.1993.1489.

9 Liisa Holm and Chris Sander. Dali: a network tool for protein structure comparison. Trends in biochemical sciences, 20(11):478–480, 1995.

10 Patrice Koehl. Protein structure similarities. Current Opinion in Structural Biology, 11(3):348–353, 2001. URL: http://www.sciencedirect.com/science/article/pii/S0959440X00002141, doi:http://dx.doi.org/10.1016/S0959-440X(00)00214-1.

11 R. Kolodny and N. Linial. Approximate protein structural alignment in polynomial time. Proc Natl Acad Sci U S A, 101(33):12201–12206, August 2004. URL: http://dx.doi.org/10.1073/pnas.0404383101, doi:10.1073/pnas.0404383101.

12 Michael Levitt and Mark Gerstein. A unified statistical framework for sequence comparison and structure comparison. Proceedings of the National Academy of Sciences, 95(11):5913–5920, May 1998. URL:http://www.pnas.org/cgi/content/abstract/95/11/5913, doi:10.1073/pnas.95.11.5913.

13 Anita Lewit-Bentley and Stéphane Réty. Ef-hand calcium-binding proteins. Current opinion in structural biology, 10(6):637–643, 2000.

14 Anthony R Means, Mark FA VanBerkum, Indrani Bagchi, Kun Ping Lu, and Colin D Rasmussen. Regulatory functions of calmodulin. Pharmacology & therapeutics, 50(2):255–270, 1991.

15 Satish Chandra Panigrahi and Asish Mukhopadhyay. An eigendecomposition method for protein structure alignment. In International Symposium on Bioinformatics Research and Applications, pages 24–37. Springer, 2014.

16 Sunil Ray. Beginners Guide To Learn Dimension Reduction Techniques. https://www.analyticsvidhya.com/blog/2015/07/dimension-reduction-methods//, 2015.

17 I. N. Shindyalov and P. E. Bourne. Protein structure alignment by incremental combinatorial extension (CE) of the optimal path. Protein Engineering, 11(9):739–747, September 1998. URL: http://dx.doi.org/10.1093/protein/11.9.739, doi:10.1093/protein/11.9.739.

18 Jinrui Xu and Yang Zhang. How significant is a protein structure similarity with TM-score = 0.5? Bioinformatics, 26(7):889–895, 2010. URL: http://bioinformatics.oxfordjournals.org/content/26/7/889.abstract, arXiv: http://bioinformatics.oxfordjournals.org/content/26/7/889.full.pdf+html, doi:10.1093/bioinformatics/btq066.

19 Yang Zhang and Jeffrey Skolnick. Scoring function for automated assessment of protein structure template quality. Proteins: Structure, Function, and Bioinformatics, 57(4):702–710, 2004. URL:http://dx.doi.org/10.1002/prot.20264, doi:10.1002/prot.20264.

